# PBMCpedia: A Harmonized PBMC scRNA-seq Database With Unified Mapping and Enhanced Celltype Annotation

**DOI:** 10.1101/2025.08.06.668843

**Authors:** Emma S. Hoffmann, Lara Hombrecher, Ian F. Diks, Matthias Flotho, Andreas Keller, Friederike Grandke

## Abstract

Reproducibility in single-cell transcriptomics remains limited by inconsistent preprocessing, heterogeneous cell type annotations, and study-specific batch effects. This challenge is particularly pronounced in peripheral blood mononuclear cell (PBMC) datasets, which are central to immunological research but rarely harmonized across studies. We present PBMCpedia, a unified PBMC atlas comprising over 4.3 million single cells from 519 samples across 24 publicly available scRNA-seq studies. Unlike prior efforts, PBMCpedia reprocesses all raw sequencing data using a single, standardized pipeline with consistent quality control, batch correction, and multi-resolution cell type annotation. The dataset spans 14 diseases, including autoimmune, infectious, and neurodegenerative conditions, alongside healthy controls, enabling reproducible, metadata-aware comparisons across biological contexts. In addition to transcriptomes, PBMCpedia includes TCR/BCR repertoire data for 75 samples and surface protein measurements for 56 samples, supporting integrative immune profiling at the transcriptomic and proteogenomic levels. To support exploration and accessibility, we provide an interactive web interface (https://web.ccb.uni-saarland.de/pbmcpedia/) for querying gene expression, marker genes, and pathway enrichment across cell types, conditions, sexes, and age groups. PBMCpedia fills a critical gap by offering a transparent, harmonized, and disease-diverse PBMC resource designed for cross-study immune profiling and discovery.

## Introduction

Single-cell RNA sequencing (scRNA-seq) of peripheral blood mononuclear cells (PBMCs) is a powerful approach to profile immune heterogeneity across diverse biological contexts, including infection, autoimmunity, aging, and neurodegeneration. Despite the availability of hundreds of public PBMC datasets, comparative analyses remain difficult due to inconsistent preprocessing, divergent annotation schemes, and batch effects. These challenges limit reproducibility and hinder the reuse of existing datasets.

Several large-scale PBMC atlases have addressed specific aspects of this issue. The Allen Institute’s Human Immune Health Atlas (1, 2) includes over 16 million immune cells from healthy donors and provides a standardized annotation framework stratified by age, but excludes disease cohorts. Jiménez-Gracia et al. (3) reprocessed raw data from 356 patients across 18 inflammatory conditions, generating the “Inflammation Landscape” using interpretable machine learning. However, their harmonized dataset is not accessible via an interactive platform or downloadable in full. Gibson et al. (4) created a PBMC aging atlas by integrating 2.8 million cells from 35 studies, but used heterogeneous preprocessing pipelines and do not provide a centralized portal. Cancer-SCEM 2.0 (5) aggregates over 1,400 datasets across 74 cancer types, with limited emphasis on PBMCs.

To address these limitations, we developed **PBMCpedia**, a harmonized, multi-disease PBMC atlas comprising 4,293,193 single-cell transcriptomes from 519 samples across 24 publicly available scRNA-seq studies (Supplementary Table 1). Unlike prior efforts, PBMCpedia uniformly reprocesses raw FASTQ files using a consistent pipeline with standardized quality control, batch correction, and hierarchical annotation. Whenever available, all samples include metadata on age, sex, and disease status, enabling stratified, demographic-aware analyses (Figure 1).

**Fig. 1.**
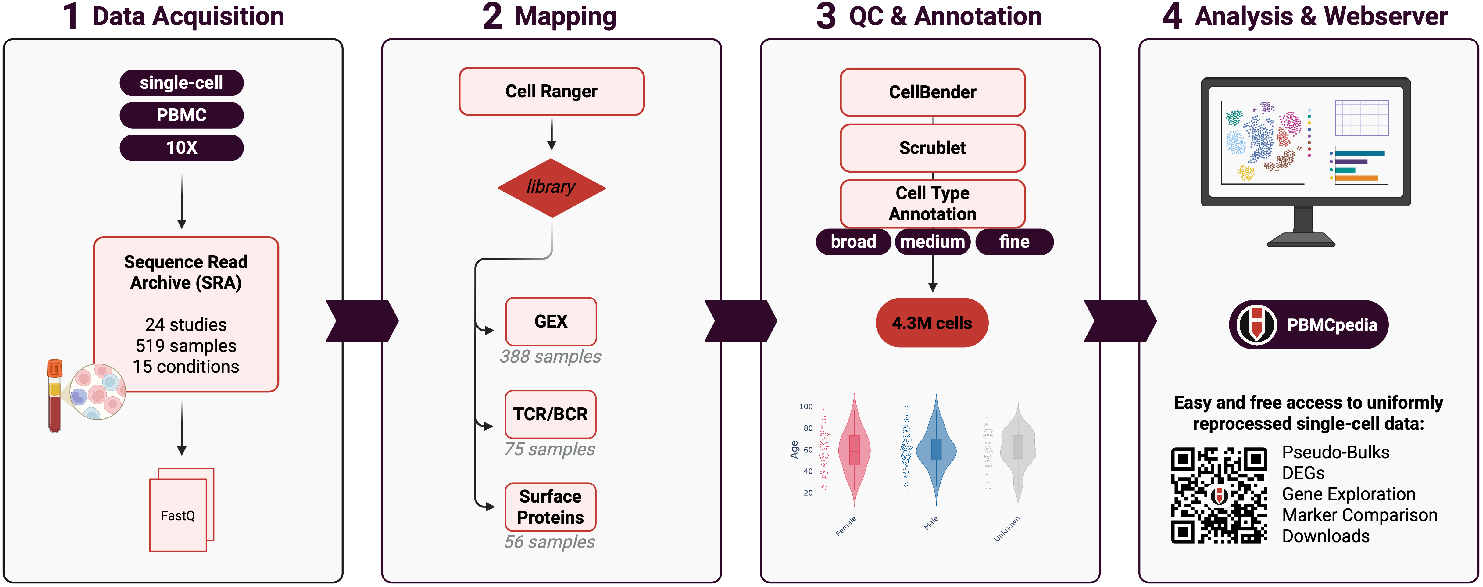
Graphical abstract.

## Materials and Methods

We included 24 publicly available scRNA-seq PBMC datasets covering 14 disease conditions and healthy controls. All datasets were generated with 10x Genomics technology, included raw FASTQ files, and contained usable sample metadata (age, sex, disease status). Datasets lacking raw data or essential metadata were excluded. In total, 519 samples and 4,293,193 high-quality cells were retained following uniform quality control.

### Data Acquisition

Sequencing data were retrieved from the Sequence Read Archive (SRA) using the SRA Toolkit (v3.0.0). We downloaded `.sra` files using `prefetch`, verified them with `vdb-validate`, and extracted paired reads using `fasterq-dump` with `-split-files` and `-include-technical` to preserve index reads. Because submission formats varied, files were manually curated to ensure correct pairing and orientation.

Metadata were collected via literature review and the NCBI SRA Run Selector. Age was treated as a continuous variable; sex was categorized as male, female, or unknown to reflect inconsistent reporting.

### Uniform Preprocessing and Quality Control

All datasets were aligned with Cell Ranger (v9.0.0) to GRCh38-2024-A. Count matrices were processed using Scanpy (v1.11.1) (6), CellBender (v0.3.2) (7) for ambient RNA correction, and Scrublet (via Scanpy) (8) for doublet detection. Cells with fewer than 200 genes or above the 98th percentile for gene/UMI counts were removed.

Integration was performed using Harmony (via harmonypy, version 0.0.10) (9) on 15 PCs, correcting batch effects by project and sample ID. UMAP visualizations confirmed dataset mixing and clustering by immune identity.

Multiplexed samples were demultiplexed by mapping cells to published donor assignments. This ensured accurate donor attribution.

### Annotation Strategy

Cell type labels were transferred using the Allen Institute’s `cell_type_mapper` tool at three hierarchical levels: AIFI_L1 (major lineages), AIFI_L2 (broad subtypes), and AIFI_L3 (fine-grained subsets) using the Human Immune Health Atlas (1) as reference. This annotation supports flexible downstream analysis.

### Differential Expression and Pathway Analysis

Pseudobulk DGE analysis was conducted using Scanpy (v1.11.2) (6) and limma (10) (v3.58.1 via rpy2 v3.5.11), comparing disease and control samples across cell types, annotation levels, sexes, and age groups. Pathway enrichment was performed using `gseGO` from clusterProfiler (version 4.10.0) (11).

For each comparison, DGE was computed using the Wilcoxon rank-sum test with Benjamini-Hochberg correction. Stratifications included sex (female/male vs. same-sex controls) and age group (young: <25; adult: 25–64; elderly: >64), with an additional pooled comparison to maximize power.

### Cross-Study Reproducibility

We evaluated reproducibility by comparing DEGs across two COVID-19 datasets ((12), (13)) before and after harmonization. Correlation of log fold changes (Pearson’s *r*), Tanimoto Coefficient (14), and overlap of significant DEGs were used to assess agreement.

Pathway concordance was assessed by semantic similarity of enriched GO terms using the Wang method, implemented in GOATOOLS (15). Redundant GO terms were reduced using `minimal_set` from ontologyIndex (16).

### Machine Learning Experiments

We tested model generalizability using incremental training of a Multi-layer Perceptron (MLP) classifier (scikit-learn v1.5.2) (17) across studies. For each of three annotation levels, we trained on sequential batches and evaluated on a held-out project (P09) using `partial_fit`. Models were tuned across hyperparameters including `alpha` (0.0001; 0.001; 0.01), and loss function (constant; invscaling; adaptive) to optimze the weighted F1 score. Accuracy, precision, recall, and F1 score were recorded after each update.

### Web Server Implementation

PBMCpedia is hosted as a Django-based web application with visualizations rendered via Plotly and DataTables.

Integrated knowledge bases, including CellMarker (18), PanglaoDB (19), MSigDB (20, 21), and the Human Protein Atlas (22) enhance interpretability. Contextual links for each gene allow real-time reference to known functions and expression profiles.

## Results

### Overview

PBMCpedia integrates 24 publicly available scRNA-seq datasets into a harmonized reference atlas of human peripheral blood mononuclear cells. After uniform reprocessing and stringent quality control, the dataset comprises 4,293,193 high-quality transcriptomes from 519 samples, covering 14 disease conditions and healthy controls (Figure 2A).

**Fig. 2.**
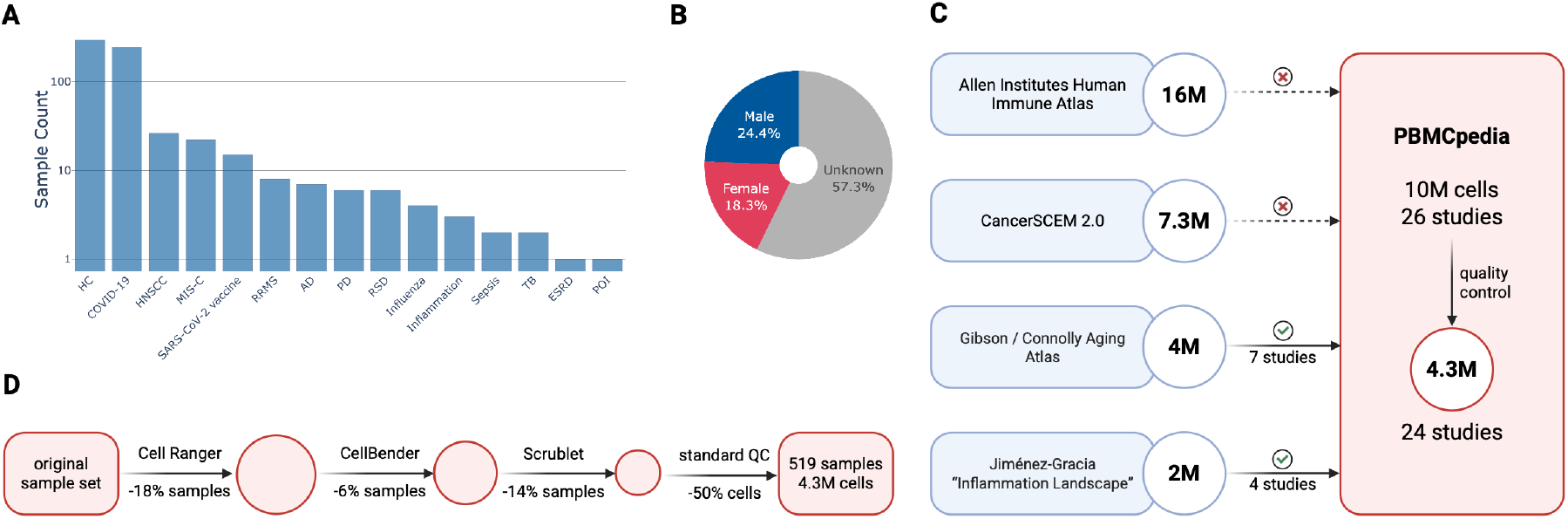
**(A)** Disease distribution **(B)** Overall sex distribution across all 519 samples. **(C)** Overlap of studies included in PBMCpedia compared to four major PBMC atlases: the Allen Immune Human Health Atlas, Jiménez-Gracia’s Inflammation Landscape, Gibson et al.’s aging atlas, and CancerSCEM 2.0. PBMCpedia shares a small subset of studies with Gibson and Jiménez-Gracia but is otherwise largely non-overlapping. **(D)** Cell and sample filtering across preprocessing steps.

The resource spans a broad range of biological contexts, including infectious diseases (e.g., COVID-19, tuberculosis), autoimmune disorders (e.g., systemic lupus erythematosus, rheumatoid arthritis), and neurodegeneration (e.g., Alzheimer’s and Parkinson’s disease). Each sample is (where available) annotated with standardized metadata including disease status, donor sex, and age, enabling stratified analysis across demographic variables (Figure 2B).

All datasets were processed through a consistent pipeline and integrated using Harmony to correct for batch effects while preserving biological structure. Cells were annotated using a three-level hierarchical framework aligned with the Allen Human Immune Health Atlas, capturing both broad immune lineages and fine-grained subtypes.

Where available, PBMCpedia also includes multimodal profiles: 75 samples contain paired TCR/BCR repertoire sequencing, and 56 include CITE-seq–based surface protein quantification. These modalities enable joint analysis of gene expression, immune clonality, and protein-level phenotypes. In total, PBMCpedia provides a reproducible, richly annotated PBMC resource suitable for comparative analysis, biomarker discovery, and immunological exploration across diverse diseases and patient subgroups.

### Comparison to Existing Resources

PBMCpedia builds on prior atlases while addressing key limitations (Figure 2C). The Allen Human Immune Health Atlas (1, 2) profiled 1.8 million PBMCs from over 100 healthy donors, later expanding to over 16 million immune cells to examine age-related changes. However, it excludes disease cohorts. In contrast, PBMCpedia spans a wide range of disease conditions. Jiménez-Gracia et al. (3) reprocessed 2 million PBMCs from 356 patients across 18 inflammatory conditions in the “Inflammation Landscape”. While harmonized, that dataset lacks a public browser and downloadable processed data. PBMCpedia offers both interactive exploration and full access to data and code. Gibson et al. (4) integrated 2.8 million cells from 35 public studies into an aging-focused PBMC atlas, supplemented by a 1.2-million-cell validation cohort. However, they used heterogeneous pipelines and do not provide a unified, interactive portal. CancerSCEM 2.0 (5) aggregates over 1,400 datasets across 74 cancer types, including PBMCs. However, its primary emphasis is on tumor microenvironments rather than systemic immune profiling.

PBMCpedia offers several advantages. All datasets are reprocessed from raw FASTQ files using a consistent pipeline, eliminating batch effects from study-specific workflows. The atlas covers 14 disease contexts and healthy controls, supports hierarchical annotations aligned with the Allen framework, and provides metadata-stratified differential expression and pathway enrichment analyses. Multimodal features (TCR/BCR and CITE-seq) enable integrative exploration of transcriptomic, clonotypic, and proteomic variation.

To ensure robust and reproducible insights, we applied strict preprocessing and quality control filters across all studies. These include thresholds for gene and UMI counts, mitochondrial read fractions, and doublet exclusion based on both transcriptomic and surface protein modalities. While these steps are essential to minimize technical artifacts and ensure biological relevance, they come at a cost: over 50% (5,625,609) of initially captured cells were discarded, resulting in a substantial shrinkage of the dataset. This rigorous curation underscores PBMCpedia’s focus on data integrity over raw cell quantity, yielding a high-confidence reference atlas suitable for downstream machine learning and systems immunology applications (Figure 2D).

All results are freely accessible through a user-friendly web interface, with downloadable data and code to ensure transparency and reuse. PBMCpedia thus fills a critical gap in the single-cell landscape by offering a reproducible, disease-aware resource for comparative immunology.

### Improved Machine Learning Performance through Diversity

We evaluated classifier generalization across diverse datasets using incremental training, where each batch corresponds to one study. The weighted F1-score was tracked throughout (Figure 3A), with individual training runs in transparent gray and a smoothed average in red. Performance improves rapidly across the initial batches, reflecting consistent signal from early datasets. The model continues to refine its predictions as more heterogeneous data is added. Notably, a few studies (particularly P02 and P03) reproducibly coincide with dips in performance. This suggests they introduce higher variance or contain edge-case cell types not well represented in earlier batches. Importantly, we cannot assume that the reference annotations are biologically perfect as some disagreements may reflect real biological ambiguity or annotation uncertainty rather than model error.

**Fig. 3.**
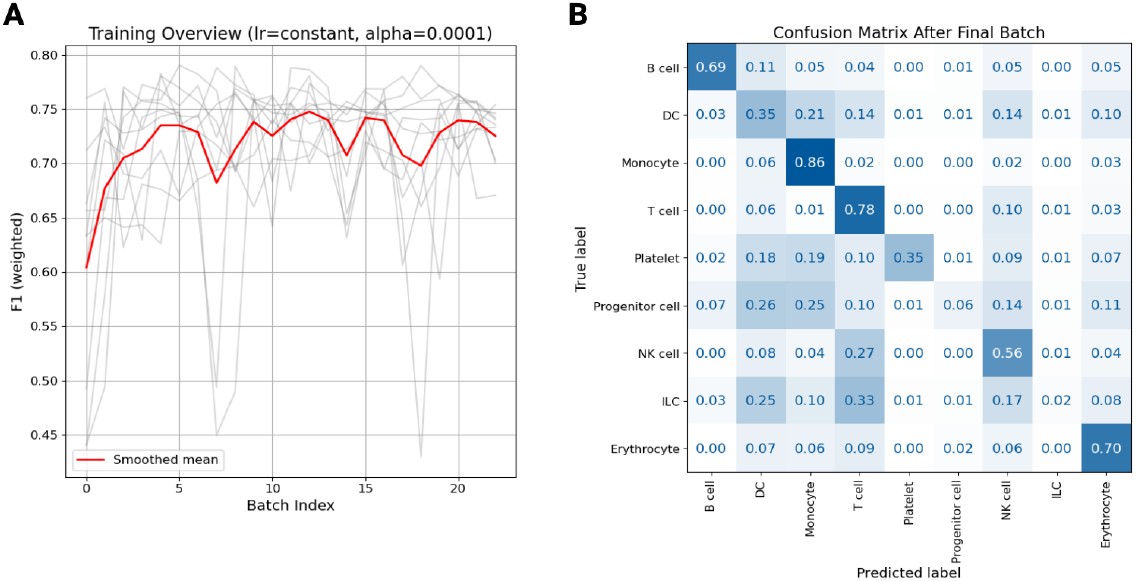
**(A)** F1-score (weighted) across training batches. Each line represents a different training run with permuted batch/project order; the red line shows the smoothed mean. **(B)** Normalized confusion matrix after the final training batch, showing predicted versus true cell type labels.

Rather than being a limitation, this behavior offers a valuable diagnostic window: the incremental setup allows us to identify where and why performance fluctuates, leading to better understanding of data quality and diversity. Importantly, despite local drops, the overall performance remains high and stable, highlighting that our training approach is robust to heterogeneity.

The final confusion matrix (Figure 3B) highlights the overall classification performance, with high diagonal values for most major cell types. Monocytes (0.86), T cells (0.78), and erythrocytes (0.70) are classified with high confidence. As expected, overlap occurs among phenotypically similar or transcriptionally related populations, such as DCs, progenitors, and ILCs, where shared marker expression may blur class boundaries.

### Cross-Disease Immune Signatures

Because PBMCpedia spans diverse biological contexts, all processed through a unified pipeline, it enables direct, systematic comparisons across conditions. Users can explore differences and commonalities in immune responses across diseases (e.g., influenza versus COVID-19), age groups (e.g., young versus adult), and sexes (e.g., male versus female) without the confounding effects of batch differences or inconsistent annotations. This level of comparability is difficult to achieve when integrating data from separate studies, where technical variation and divergent labeling schemes often obscure true biological signals.

With that, PBMCpedia supports demographic filtering, allowing users to restrict analyses to subsets such as female patients over 60. Even within such filters, sample sizes remain large enough to detect meaningful differences, facilitating age- and sex-aware disease comparisons.

## Discussion

PBMCpedia enables robust, comparative immune profiling across diseases by offering a harmonized and openly accessible dataset with standardized preprocessing. Its three-level annotation system serves both novice users and expert immunologists, providing flexibility for various analytical depths. Stratified analyses by sex and age, together with multimodal integration where available (TCR/BCR, CITE-seq), further enhance its utility for exploring immune heterogeneity across biological and clinical contexts.

As a community-curated resource, PBMCpedia reflects the availability of public data. Some diseases, including cancers and rare conditions, remain underrepresented. Future versions will incorporate new studies as they become available, leveraging our reproducible pipeline to expand coverage. This includes especially the raw data of the Allen Immune, facilitated by the already shared annotation, and the “Inflammation Landscape” (3).

While transcriptomics is the core modality, PBMCpedia already includes some multimodal datasets. Future updates will expand this further, including spatial transcriptomics and cell-cell interaction networks. Despite robust batch correction, residual cohort differences may persist. We encourage users to leverage the provided metadata, especially age, sex, and control samples from multiple studies, for careful interpretation.

PBMCpedia addresses the growing need for reproducible, cross-disease immune profiling by enabling scalable, metadata-aware analysis and therefore supporting both discovery and hypothesis-driven research. All data, code, and results are freely available through an interactive web platform.

By prioritizing accessibility, consistency, and scalability, PBMCpedia serves as a valuable tool for immunologists, computational biologists, and clinical researchers. We invite community feedback and contributions to expand and refine PBMCpedia as an open resource for immune research.

## Acknowledgements

The authors used ChatGPT (OpenAI, GPT-4o) to assist with phrasing and language refinement during manuscript preparation. All scientific content, data analysis, figure creation, and the initial manuscript draft were carried out by the authors, who also thoroughly reviewed and revised all text. Figures were created using BioRender.com. The project was supported by the Michael J. Fox Foundation for Parkinson’s Research [MJFF-021418 to A.K. and T.W-C., 14446, 17047] the Schaller-Nikolich Foundation (to A.K.); Saarland University. Computational resources used within this study were financed through the DFG [466168626 to A.K.].

## Data Availability

PBMCpedia is freely available at https://web.ccb.uni-saarland.de/pbmcpedia/.

---

**Supplementary Table 1.** Each row corresponds to one of the 24 publicly available single-cell PBMC studies integrated into PBMCpedia. For each study, we list the internal project ID used throughout the resource, the associated disease(s), the number of high-quality cells retained after preprocessing, and the modalities available. Modalities include gene expression (GEX), TCR/BCR repertoire profiling, and surface protein quantification (CITE-seq).

| ID | Study | Diseases | # Cells | Modalities |
| --- | --- | --- | --- | --- |
| P01 | Luo et al. (2022) (23) | - | 139,111 | GEX, TCR/BCR, surface protein |
| P02 | Smith et al. (2020) (24) | - | 9,620 | GEX |
| P03 | Yang et al. (2019) (25) | - | 8,369 | GEX |
| P04 | Yazar et al. (2022) (26) | - | 1,539,942 | GEX |
| P05 | Volden et al. (2022) (27) | - | 3,131 | GEX |
| P06 | Goudot et al. (2017) (28) | - | 850 | GEX |
| P07 | Xu and Jia (2021) (29) | Alzheimer's Disease | 20,671 | GEX |
| P08 | Xiong et al. (2021) (30) | Alzheimer's Disease | 74,109 | GEX |
| P09 | Bai et. al (2022) (31) | COVID-19 | 211,272 | GEX |
| P10 | Ivanova et al. (2023) (12) | COVID-19 | 507,223 | GEX, TCR/BCR, surface protein |
| P11 | Arunachalam et al. (2020) (32) | COVID-19 | 60,655 | GEX |
| P12 | Van der Wijst et al. (2021) (13) | COVID-19 | 25,848 | GEX, surface protein |
| P13 | Liu et al. (2021) (33) | COVID-19 | 1,047,036 | GEX, TCR/BCR, surface protein |
| P14 | Ramaswamy et al. (2021) (34) | COVID-19 | 252,643 | GEX |
| P15 | Unterman et al. (2022) (35) | COVID-19 | 95,819 | GEX |
| P16 | Lee et al. (2020) (36) | COVID-19, flu | 96,992 | GEX |
| P17 | Zhang et al. (2023) (37) | End Stage Renal Disease | 9,027 | GEX |
| P18 | Cillo. (2019) (38) | Head and Neck Squamous Cell Carcinoma | 61,396 | GEX |
| P19 | Zhou et al. (2020) (39) | Inflammation | 13,933 | GEX |
| P20 | Seyedsadr et al. (2023) (40) | Multiple Sclerosis | 28,970 | GEX |
| P21 | Wang et al. (2022) (41) | Parkinson's Disease | 27,427 | GEX |
| P22 | Zhang et al. (2023) (42) | Premature Ovarian Insufficiency | 730 | GEX |
| P23 | Qiu et al. (2021) (43) | Sepsis | 17,364 | GEX |
| P24 | Cai et al. (2020) (44) | Tuberculosis | 41,055 | GEX |

## Notes

https://web.ccb.uni-saarland.de/pbmcpedia/

### Competing Interest Statement

The authors have declared no competing interest.

